# Structural Evidence for Occupancy-Dependent Inter-Site Coupling in Human Glutathione Synthetase

**DOI:** 10.64898/2026.06.09.730884

**Authors:** Matthew Stankus, Mary Anderson

## Abstract

Human glutathione synthetase (hGS) is a negatively cooperative ATP-grasp enzyme that catalyzes the final step in the biosynthesis of glutathione, a tripeptide antioxidant critical for life. hGS functions as an obligate homodimer with one active site per subunit; the two active sites are separated by ∼40 Angstroms. How ligand binding in one subunit reshapes the distant partner active site has remained a central unresolved question in understanding hGS regulation. This study provides the first atomistic model of ligand-dependent inter-subunit communication underlying negative cooperativity in hGS. Using atomistic simulations and dynamical network analysis, this study reveals how reactant- and product-bound states remodel the empty partner active site, redistribute inter-subunit interactions, and organize long-range communication between the two active sites. The product-bound/partner-empty state displayed a larger and less hydrated empty active site, demonstrating that ligand identity in one subunit alters both the geometry and solvent environment of the opposite site. Changes in ligand-dependent interactions are distributed across the dimer interface, with prominent contributions from the 42-46 interface region, the 11-30 region, and the 212-236 helical/interface region. Suboptimal path analysis shows product- and reactant-bound states share a communication scaffold, with 64.1% of transmission residues common to both pathways, 30.8% product-specific, and 5.1% reactant-specific. Together, the present results establish a detailed structural framework for hGS negative cooperativity in which ligand binding remodels a distributed allosteric network linking substrate-binding loops, the dimer interface, and the partner active site. More broadly, this work demonstrates how atomistic simulations can resolve long-range active-site coupling in multimeric enzymes and provides a foundation for experimental tests of allosteric transmission in hGS.

## 1. Introduction

Glutathione synthetase catalyzes the second and final step in the biosynthesis of glutathione, the ATP-dependent ligation of glycine to γ-glutamylcysteine to form the antioxidant tripeptide glutathione [1]. Glutathione is present in most bacterial, plant, and mammalian cells and is required for protection against reactive oxygen species, detoxification of damaging compounds, maintenance of reduced protein thiols, and regulation of the cellular redox state [2]. Decreased glutathione levels are associated with a variety of pathological conditions, and inherited deficiencies in glutathione synthetase can result in metabolic acidosis, hemolytic anemia, neurological disorders, and premature death [3]. Thus, improved understanding of human glutathione synthetase is important not only for defining the molecular basis of glutathione biosynthesis, but also for understanding how changes in enzyme structure and regulation may contribute to disease.

Human glutathione synthetase (hGS) is a member of the ATP-grasp superfamily of enzymes, which use ATP to drive carbon-nitrogen bond formation through an acyl-phosphate intermediate [4]. Unlike many structurally characterized ATP-grasp enzymes, hGS is a mammalian homodimer and displays negative cooperativity toward its γ-glutamyl substrate [5] [6]. In negatively cooperative enzymes, the binding of substrate to one active site decreases the affinity of a distant active site for the same substrate. This type of regulation is less well understood than positive cooperativity. Still, it plays a critical role for branch-point enzymes that control metabolite flux over a wide range of substrate concentrations. From an evolutionary perspective, negative cooperativity allows multimeric enzymes to avoid abrupt saturation, tune metabolic flux in response to changing substrate concentrations, and maintain reserve catalytic capacity under metabolic stress. In hGS, negative cooperativity may help regulate γ-glutamylcysteine utilization while maintaining cellular glutathione levels and preserving heightened capacity for glutathione biosynthesis.

Previous kinetic studies using the γ-glutamylcysteine analog γ-glutamyl-α-aminobutyrate (GAB) report a Hill coefficient of ∼0.69, indicating hGS displays negative cooperativity towards the γ-glutamyl substrate [5]. GAB is used as a non-thiol analog of γ-glutamylcysteine because it maintains steric similarity and comparable activity while avoiding cysteine thiol oxidation, thereby simplifying assay experiments [5] [7]. Thus, this congener preserves the relevant substrate features needed to probe active-site organization and inter-subunit communication.

Prior attempts to experimentally isolate a monomeric, independently folded, and catalytically competent subunit have been unsuccessful, thereby limiting traditional methods for studying asymmetric ligand binding in hGS [5]. Thus, the structural basis of hGS negative cooperativity remains poorly defined. The enzyme is an obligate homodimer with two active sites separated by ∼40 Å and connected through a relatively small dimer interface [6]. Thus, substrate binding at one active site must communicate over a long distance to alter the binding properties of the partner active site. Previous experimental and computational studies identify several regions that may participate in this communication, including the dimer interface and the conserved S-, G-, and A-loops surrounding the active site [4] [8] [9] [10] [11] [12]. These studies establish that hGS allostery likely arises from communication among substrate-binding loops, catalytic loops, and inter-subunit contacts.

Prior work on hGS’s dimer interface shows that interface residues can contribute differently to stability and allostery. Hydrophobic residues Val44 and Val45 are located near the interface and are implicated in the allosteric pathway of hGS [12]. Mutations at these residues decrease thermal stability and alter cooperativity, with Val45 mutations generally producing larger effects than Val44 mutations [12]. In contrast, strong electrostatic interactions involving Asp24, Ser42, and Arg221 are essential for enzyme activity and stability, but alanine mutations at these positions do not substantially alter the Hill coefficient [10]. These prior findings indicate that strong inter-subunit electrostatic contacts help maintain the hGS dimer, whereas weaker hydrophobic and distributed interface interactions may be more directly involved in transmitting allosteric communication.

hGS’s active-site loops also play important roles in catalysis and regulation. The S-loop forms part of the γ-glutamylcysteine binding site and contains residues that directly bind or position the negatively cooperative substrate [4] [8]. Mutations at S-loop residues such as Arg267, Tyr270, and Pro272 severely reduce activity, alter substrate binding, and, in some cases, diminish negative cooperativity [8]. The G-loop glycine triad, particularly Gly369 and Gly370, is essential for ligand binding, positioning of ATP’s γ-phosphate, and active-site closure [11]. The A-loop residue Asp458 is likewise required for normal catalysis and allosteric behavior [9]. Mutation of Asp458 alters substrate binding and eliminates negative cooperativity without affecting overall hGS stability [9]. Together, these prior studies suggest that hGS allostery begins with specific substrate-loop interactions and is then transmitted through a larger structural network.

Although these studies have identified residues and loops important for hGS function, they do not attempt to explain how ligand identity in one subunit remodels the empty active site of the partner subunit. In particular, it remains unclear whether product- and reactant-bound states communicate via distinct allosteric routes or a shared pathway modulated by ligand states. It is also unknown how changes at the occupied site are structurally expressed in the empty partner site. Addressing these questions requires analysis of asymmetric occupancy states that more directly model inter-site communication within the hGS dimer.

No prior computational study has examined negative cooperativity in hGS at the current level of atomistic detail across asymmetric ligand-occupancy states while integrating empty-site remodeling, dimer-interface rewiring, and dynamic communication pathway analysis. In the present study, molecular dynamics simulations and dynamical network analysis are used to examine ligand-dependent communication between hGS subunits.

## 2. Methods

### 2.1 Molecular Dynamics System Preparation

This study analyzes wild-type human glutathione synthetase as a homodimer in ligand-occupancy states designed to isolate the structural consequences of active-site occupancy. The analyzed singly occupied states include product-bound/partner-empty and reactant-bound/partner-empty systems. Product-bound active sites contain glutathione, ADP, inorganic phosphate, and Mg^2+^. Reactant-bound active sites contain γ-glutamylcysteine, ATP, and Mg^2+^. The empty partner active sites contain no ligand, nucleotide, phosphate, or Mg^2+^.

The study uses the human glutathione synthetase crystal structure 2HGS from the RCSB Protein Data Bank as the starting structure [6]. CHARMM-GUI Solution Builder prepares the systems, and CHARMM-GUI FF-Converter converts them to AMBER-family GROMACS topologies [13] [14] [15] [16]. The simulations model the protein with the AMBER ff14SBonlysc force field, describe ligands and cofactors with GAFF2 parameters, and represent water with the TIP3P model [17]. The preparation assigns protonation states consistent with physiological pH 7.4.

Each system uses a cubic periodic box with a 10 Å solvent buffer. The preparation workflow neutralizes each system and then ionizes it to a final salt concentration of 0.15 M. Steepest-descent minimization reduces the maximum force to below 1000 kJ mol^−1^ nm^−1^. The equilibration protocol then propagates each system for 125 ps at 310.15 K with a 1 fs timestep and positional restraints on the protein backbone and side chains. Production simulations use a 2 fs timestep, LINCS constraints on bonds involving hydrogen atoms, V-rescale temperature coupling at 310.15 K, isotropic C-rescale pressure coupling at 1 bar, 0.9 nm real-space cutoffs for van der Waals and short-range Coulomb interactions, and particle-mesh Ewald electrostatics. The simulations save coordinates every 100 ps. The MD study generates three independent replicates for each occupancy state. Each replicate contains 10 continuous 100 ns production segments, yielding a nominal production length of 1.0 μs per replicate and a total aggregate production sampling of 6 μs.

### 2.2 Equilibration assessment and analysis window selection

The study restricts all analyses to accepted post-equilibration windows. This procedure reduces the influence of early relaxation and ensures that comparisons between occupancy states use stable trajectory intervals. A ligand-pocket distance state model assigns bound and unbound ligand states. The model classifies a site as bound when the ligand-pocket minimum heavy-atom distance is less than or equal to 5.0 Å, and as unbound when this distance is greater than or equal to 5.5 Å. Distances between 5.0 Å and 5.5 Å retain their previous state assignment to prevent rapid threshold switching.

Python implementation of the multistate Bennett acceptance ratio (PyMBAR) time-series methods assess equilibration and statistical inefficiency [18] [19]. For each condition, the analysis selects accepted windows after evaluating ligand-state persistence, equilibration diagnostics, and trajectory stationarity. Subsequent pocket, hydration, contact, and network analyses use these accepted windows.

### 2.3 Empty-site pocket volume and hydration

The analysis measures pockets using a predefined empty-site reference pocket rather than redefining the pocket independently in each trajectory. Local backbone alignment of the empty-site neighborhood aligns the same structural region consistently across conditions before pocket analysis.

The workflow obtains pocket volume from MDPocket descriptors after alignment to the reference empty-site pocket [20]. The analysis quantifies empty-site hydration as the number of water oxygen atoms within 6.0 Å of the aligned template-site heavy atoms. These measurements assess whether the identity of the ligand bound in one subunit alters the volume and solvent environment of the unoccupied partner active site. The study summarizes pocket volume and hydration values in 10 ns blocks before making condition-level comparisons [21] [22].

### 2.4 Inter-subunit contact and interface rewiring analysis

The analysis quantifies dimer-interface remodeling from residue-pair heavy-atom contact occupancies across the two subunits. The workflow classifies a residue pair as contacting in each frame when any non-hydrogen atom from one residue falls within 4.5 Å of any non-hydrogen atom from the opposing-chain residue. Contact occupancy represents the fraction of analyzed frames in which the residue pair satisfies this criterion.

The study evaluates ligand-dependent interface rewiring by comparing inter-chain contact occupancies between the product-bound/partner-empty and reactant-bound/partner-empty states. For each residue, the analysis sums the magnitudes of pairwise contact-occupancy differences with all opposite-chain partners to obtain a residue-level rewiring score. The workflow then normalizes these scores for structural visualization. High scores, therefore, identify residues whose inter-subunit packing responds most strongly to the identity of the ligand bound in the opposite active site. This analysis distinguishes between a model in which communication depends on a single dominant inter-subunit contact and a model in which ligand identity redistributes many weaker contacts across the dimer interface.

### 2.5 Residue interaction network and pathway analysis

The present study examines active-site communication using residue interaction networks constructed from accepted MD windows [23] [24]. These networks treat residues as nodes and persistent inter-residue contacts as edges. The analysis assigns edge weights so that residue pairs with stronger correlated motion represent more favorable communication routes [25].

The weighted implementation of suboptimal paths analysis identifies ensembles of favorable communication routes rather than a single shortest path [25]. The analysis scores residues according to their recurrence in the resulting path ensemble. High-scoring residues, therefore, represent positions that repeatedly appear as candidate relay points between the occupied and empty active sites. The study then compares shared and ligand-specific residues between product-bound and reactant-bound singly occupied states to determine whether the two states use distinct communication routes or modulate a common scaffold.

### 2.6 Software, visualization, and reproducibility

The study performs simulations and analyses using GROMACS 2025.2 [26] [27], MDAnalysis 2.10.0 [28] [29], PyMBAR 4.2.0 [19], NetworkX 3.6.1 [30], AlloViz 1.0 [31], WISP [25], MDPocket [20], and ChimeraX 1.11.1 [32] codes. The authors prepare structural figures from representative hGS conformations with mapped residue-level scores or network-derived pathways. Analysis scripts, simulation-derived outputs, and figure source data are available upon request.

## 3. Results

### 3.1 Partner occupancy reshapes the empty active site

Human glutathione synthetase (hGS) is an obligate homodimer comprised of two identical subunits, each containing a single active site [6]. Purified hGS exhibits negative cooperativity towards the reactant γ-glutamylcysteine in enzymatic assays, suggesting the ligand bound at one active site influences the partner site’s structure [5]. These subunits are arranged with C2 symmetry, meaning that one subunit can be superimposed onto the other by a 180-degree rotation around a central point [6]. In hGS, this point lies at the center of the subunit interface, represented by the yellow dashed line labeled “dimer interface” in **Figures 1A** and **1B**.

**Fig. 1:**
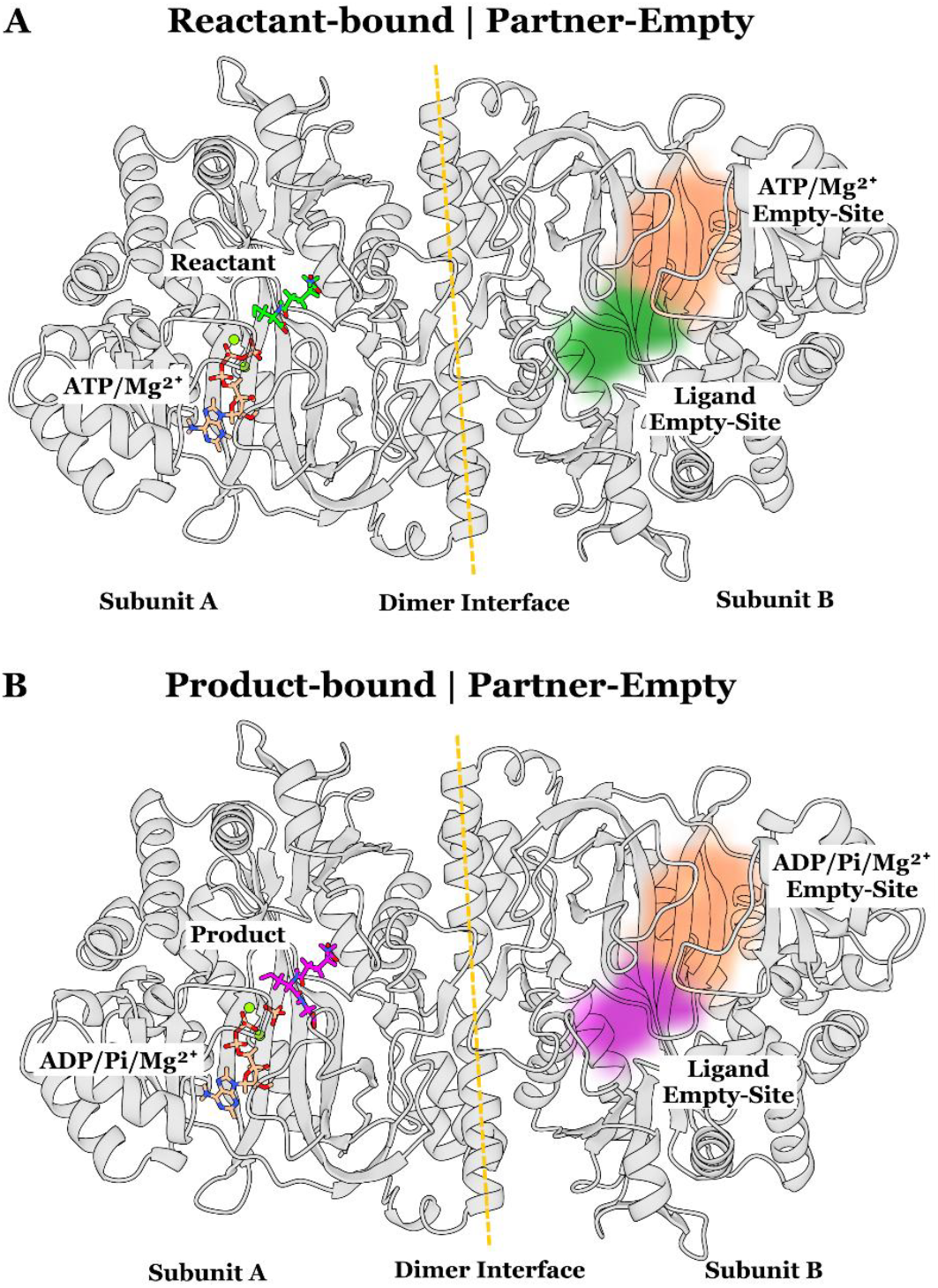
Simulation conditions used to isolate inter-site coupling effects. In both states, one subunit of the hGS homodimer is occupied, while the partner subunit is left empty. In the reactant - bound, partner-empty state (top), subunit A contains reactant and cofactors (γ-glutamylcysteine, ATP, and Mg^2+^), while subunit B contains neither ligand nor cofactors. In the product-bound, partner-empty state (bottom), subunit A contains product and cofactors (glutathione, ADP/Pi, and Mg^2+^), while subunit B likewise remains empty. The dashed line marks the dimer interface, which separates the occupied and empty-partner subunits. Colored overlays on the empty-partner subunit indicate the reference positions corresponding to where ligand and cofactor species would reside if present and are shown only to orient the reader to the empty-site geometry; they do not indicate occupancy in the partner-empty subunit.

The following analyses compare two distinct singly occupied catalytic states, as shown in **Figure 1**. The first condition, shown in **Figure 1A**, is called the reactant-bound/partner-empty state. This state represents a pre-catalytic configuration in which one subunit’s active site contains the reactant γ-glutamylcysteine, ATP, and two Mg^2+^ cofactors, while the partner subunit’s active site remains unoccupied by ligands, nucleotides, or cofactors. The second condition, shown in **Figure 1B**, is referred to as the product-bound/partner-empty state. This state represents a post-catalytic configuration in which the occupied active site contains the product glutathione, the corresponding hydrolyzed nucleotide products, inorganic phosphate, and cofactors. As in the first condition, the partner active site remains empty.

Although both subunits are catalytically functional *in vivo*, the following *in silico* analyses focus specifically on these singly occupied states. This approach allows ligand-dependent effects to be isolated and makes it possible to examine whether the ligand bound in one subunit, reactant-bound or product-bound, can influence the structure of the empty partner subunit through inter-site coupling. The initial analyses examine whether the simulated ensembles contain structural changes consistent with experimentally observed negative cooperativity.

The first analysis uses MDPocket to measure the volume of the empty partner active site (**Fig. 2A**) [20]. This method estimates the pocket volume by packing spheres into the empty site and summing the volumes of the spheres present at each time point [20]. The product-bound state has a mean empty-site volume of 1190.3 Å^3^, as indicated by the thick black line. The reactant-bound state has a lower mean empty site volume of 897.5 Å^3^. Relative to the reactant-bound/partner-empty state, the empty-site volume increases by 32.6% in the product-bound/partner-empty state. This difference corresponds to a more expanded empty-partner active-site geometry in the product-bound/partner-empty state.

**Fig. 2:**
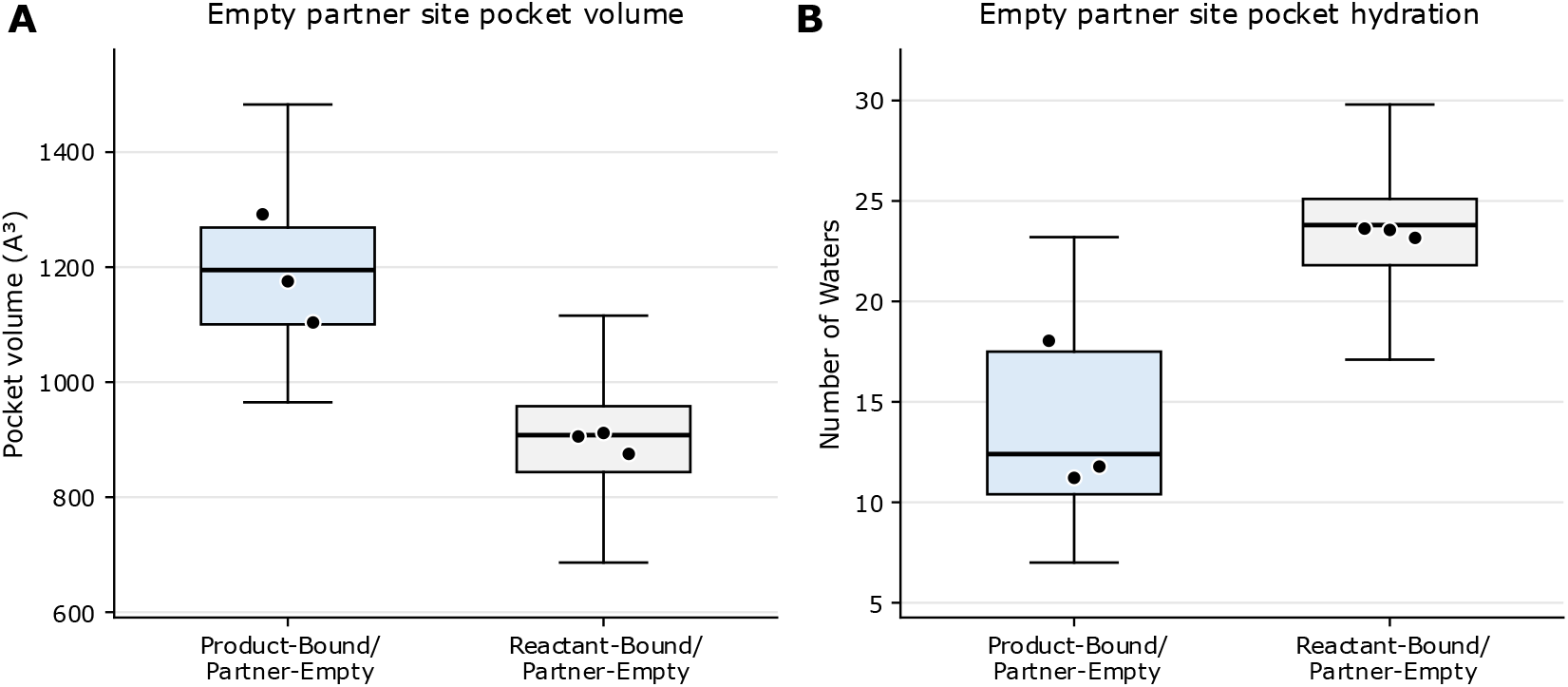
Bound-ligand identity reshapes empty-site volume and hydration. (A) Empty-site pocket volume comparing product-bound/partner-empty and reactant-bound/partner-empty states after alignment; reported pocket volume is from MDPocket descriptors. The product-bound condition showed a larger empty-site pocket volume of 32.6% (+292.8 Å3; 95% paired bootstrap CI [228.5 Å^3^, 386.3 Å^3^], n=3). (B) Empty-site hydration comparing product-bound/partner-empty and reactant-bound/partner-empty states using water-oxygen counts within the MDPocket-defined pocket volume. The product-bound condition showed that the empty site is less hydrated by 41.7% (i.e., -9.8 waters; 95% paired bootstrap CI: [-12.3, -5.6], n=3).

The second analysis evaluates empty site hydration by counting water molecules within the same region defined in the volume analysis. In most instances, one would expect a larger pocket volume to also show an increase in hydration. Counterintuitively, hydration shows the opposite trend (**Fig. 2**): despite its larger empty-site volume, the product-bound condition on average contains fewer waters than the reactant-bound condition, with means of 13.7 and 23.4 waters, respectively. Relative to the reactant-bound state, the product-bound state has an empty site that is 41.7% less hydrated.

### 3.2 Dimer interface rewiring is driven by ligand identity

Negative cooperativity depends on physical interactions between active sites for communication. For the ligand bound at one active site to influence the partner site, the effect must first propagate through the occupied subunit, towards and then across the subunit-subunit boundary, and then enter the partner subunit. The dimer interface is the structural region where communication crosses from one subunit to the other. Inter-site communication would therefore depend on either A) a narrow route through a small number of dominant residue-residue interactions, or B) through a broader redistribution of inter-subunit interactions across the interface.

Residue–residue interaction frequencies across the dimer interface are used to distinguish these communication models. A residue pair is considered interacting if any non-hydrogen atom from one residue is within 4.5 Å of any non-hydrogen atom from the opposing residue. Interaction frequencies are compared between the product-bound/partner-empty and reactant-bound/partner-empty states to isolate ligand-dependent changes from the baseline dynamics of the dimer interface. Pairwise differences are aggregated by residue and normalized to a scale from 0 to 1, producing an interface interaction score. Higher scores indicate greater ligand-dependent changes in inter-subunit interactions. A serial route is characterized by one or a few localized residues with high interaction scores, whereas distributed coupling is indicated by many residues across the interface exhibiting high interaction scores.

These interaction scores are mapped onto the corresponding residues as a color gradient (**Fig. 3**), revealing ligand-dependent interface changes are distributed across the dimer interface rather than localized to a single dominant contact. One prominent region occurs in the 42–46 loop, where 42-SEVVS-46 in the bound subunit contacts the corresponding 42-SVVES-46 segment in the empty partner subunit in an antiparallel, homotypic arrangement. This segment contains polar serine residues, an acidic glutamate, and hydrophobic valines, creating a chemically mixed interface in which local packing and polar contacts are positioned within the same short loop, while being interface accessible. A second high-scoring region appears within the 212–236^†^ α-helix of the bound subunit, which contacts the 11–30 α-helix of the empty partner subunit. These α-helical regions display a similar pattern of antiparallel symmetry, as the occupied subunit’s 11-30 α-helix region also shows high interaction scores with the empty subunit’s 212-236 α-helical region.

**Fig. 3:**
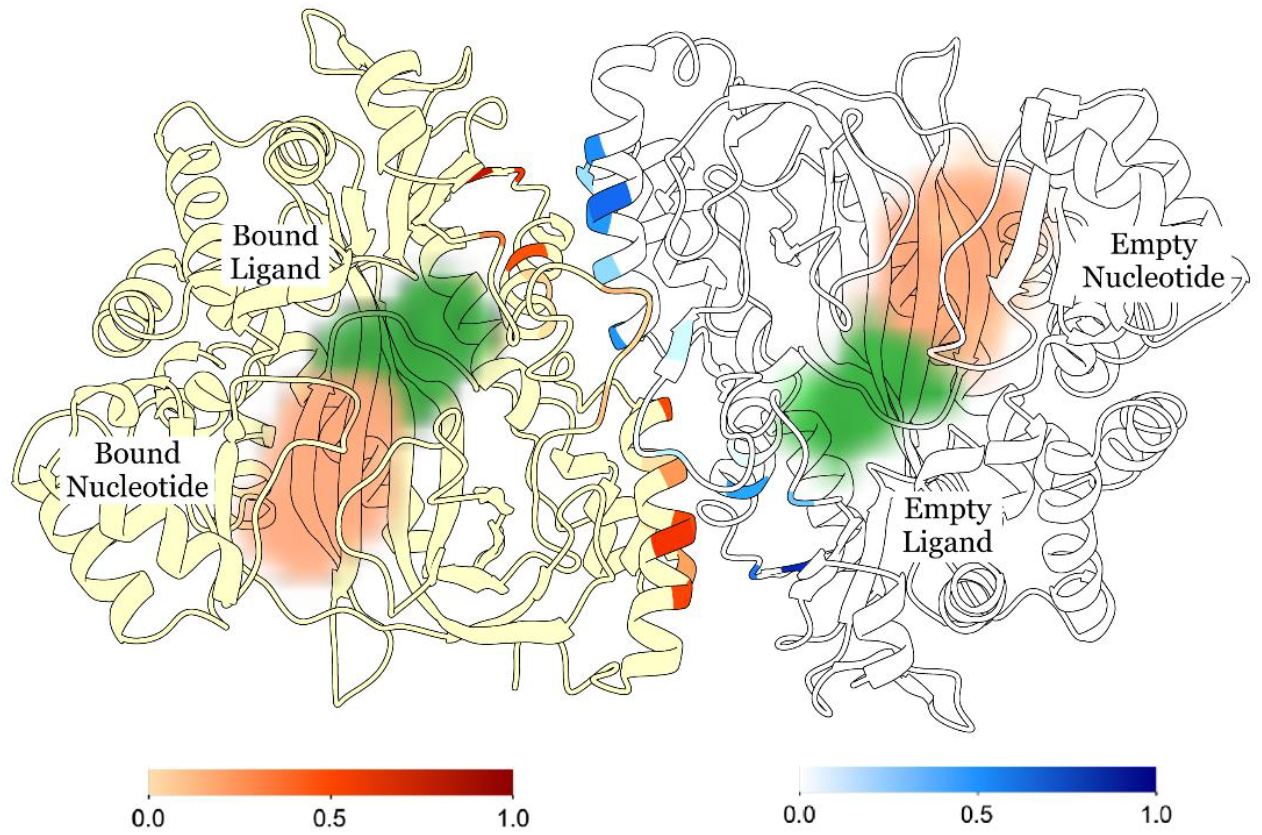
Bound-ligand identity rewires dimer interface. Residue-level interface rewiring scores for the product-bound/partner-empty versus reactant-bound/partner-empty comparison mapped onto the hGS homodimer. Inter-subunit heavy-atom contacts are quantified for all residue pairs spanning the two chains, and pairwise contact-occupancy differences between states are aggregated into per-residue rewiring scores, defined as the summed magnitude of contact-occupancy differences across opposite-chain interaction partners. Higher scores indicate residues that contribute more strongly to ligand-dependent redistribution of inter-subunit packing, with scores for the bound subunit shown in red and scores for the empty subunit shown in blue. Bound ligand and bound nucleotide are shown in the occupied subunit, and the corresponding empty ligand and nucleotide sites are labeled in the partner subunit. The broad spatial distribution of high-scoring residues argues against a single dominant serial contact pathway and instead supports distributed coupling across the dimer interface.

### 3.3 Network analyses identify a shared mediator scaffold

The foregoing analyses indicate that subunit-subunit communication is transmitted across a broad spectrum of dimer interface residues. We now analyze the hGS dimer interface and binding regions to assess how substrate binding at one subunit initiates structural reorganization of the unbound subunit. In many cooperative proteins, allosteric pathways typically emerge from multiple minor effects distributed across partially redundant, parallel routes [25] [23]. Thus, for hGS, an important unresolved question is whether product- and reactant-bound states communicate between substrate-binding regions through a common residue interaction corridor or through distinct ligand-dependent corridors.

The weighted implementation of the sub-optimal paths (WISP) method identifies candidate communication corridors through dynamic network analysis of the MD trajectories [25]. In this method, the enzyme is represented as a residue interaction network, in which residues are treated as nodes and edges connect proximal residue pairs that exhibit correlated motion [25] [23]. Edges are assigned weights, so strongly correlated residue pairs are considered as more favorable communication routes in the network. WISP then calculates the shortest optimal path(s) and an ensemble of near-optimal paths between defined source residues in the bound active site and sink residues in the unbound active site.

WISP analysis indicates transmission from the occupied active site to the empty partner site occurs through an allosteric communication network that is largely conserved between ligand states. Across the product- and reactant-bound ensembles, 64.1% of the identified transmission residues are shared, suggesting that both states use a common site-to-site communication corridor rather than fully distinct pathways. Interestingly, pathway membership differs between states: a larger proportion of residues is unique to the product-bound pathway (30.8% of all transmission residues), whereas only 5.1% are unique to the reactant-bound condition. When these residue interactions are mapped to the hGS structure (**Fig. 4**), the conserved communication route becomes apparent. The shared interactions, shown in purple (**Fig. 4A**), define the dominant transmission corridor linking the bound and empty sites. Product-bound specific interactions (**Fig. 4B**), shown in blue, remain clustered along this central route, whereas reactant-bound-specific interactions (**Fig. 4C**), shown in red, converge on the same route but also extend around the occupied active site, forming an additional active-site-proximal network. Thus, ligand identity does not recruit entirely distinct communication pathways. Instead, the product- and reactant-bound states modulate a common allosteric scaffold through ligand-specific variations in pathway membership.

**Fig. 4.**
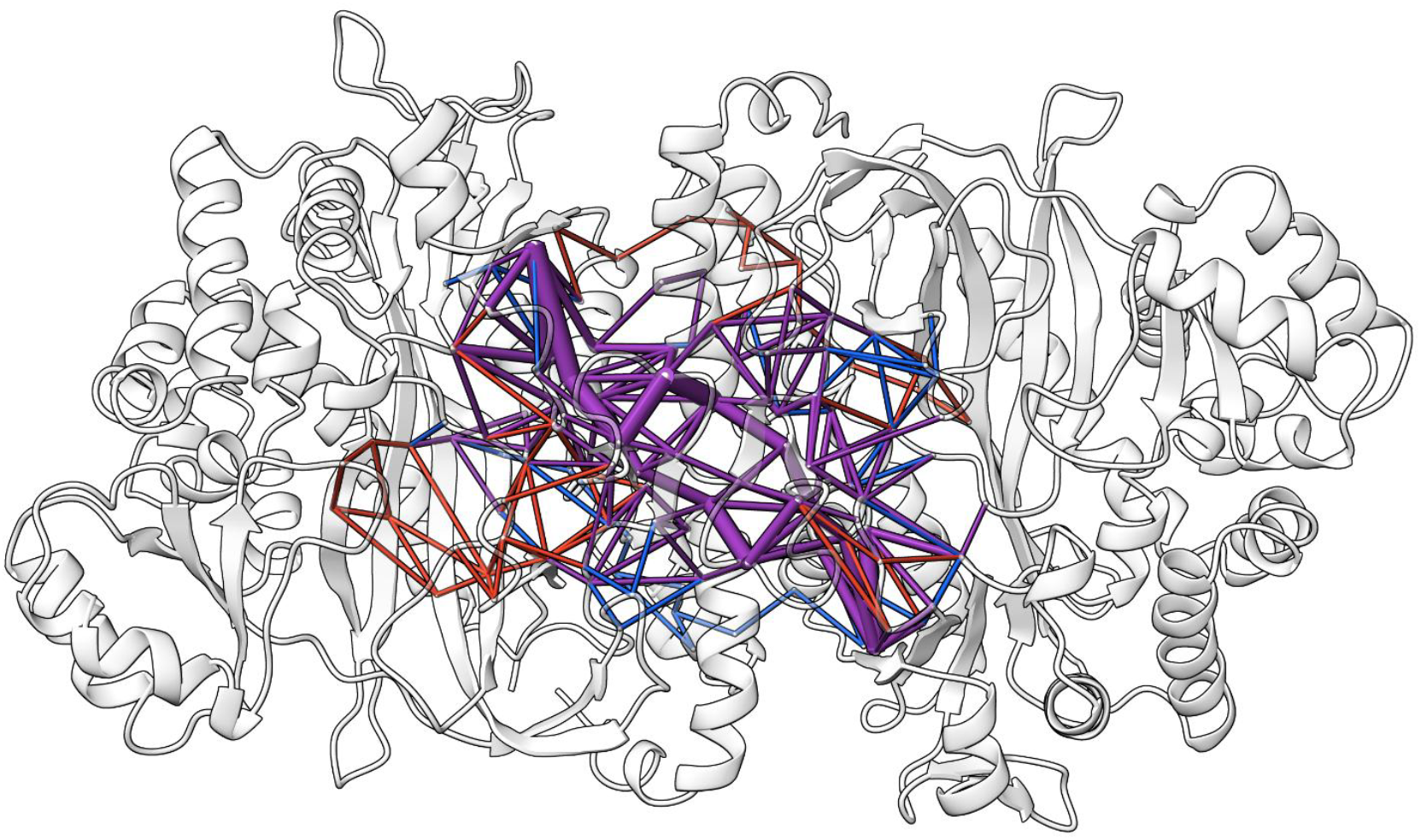
Recurrent mediator scaffold linking active sites across the dimer interface. The shared interactions shown in purple define the dominant transmission corridor linking the bound and empty sites. Product-bound specific interactions shown in blue remain clustered along the central route. Reactant-bound-specific interactions shown in red converge on the same central route but also extend around the occupied active site.

## 4. Discussion

This study supports the conclusion that ligand occupancy in one subunit of human glutathione synthetase is communicated to the partner subunit and is associated with a distinct structural state of the unoccupied partner active site. Since hGS is an obligate homodimer whose active sites are separated by ∼40 Å, any communication between those distant sites must involve a structural link to facilitate transmission. Present findings show the empty pocket becomes larger but less hydrated in the product-bound condition, suggesting the empty partner site adopts a ligand-dependent conformational and hydration state, supporting a model in which negative cooperativity reflects a shift in the conformational ensemble of the unoccupied active site rather than a purely local binding-site effect.

Broadly, the present results support a model of enzyme allostery in which communication between distant active sites results in ligand-dependent remodeling of partner sites. In this model, negative cooperativity does not require a single dominant allosteric contact or a radical global conformational change. Instead, ligand binding can influence active-site hydration, pocket geometry, and loop organization by altering interactions within a conserved communication scaffold. This framework may be relevant to other oligomeric enzymes in which active sites are physically separated but functionally coupled [23], especially when mutations outside functional hotspots alter activity, stability, or cooperativity [8]. Thus, hGS provides a useful example of how distributed communication networks can link substrate binding at one active site to altered readiness of distant active sites.

The present results build on earlier work concerning the hGS dimer interface. Prior mutagenesis and molecular dynamics studies show that Val44 and Val45 are important for hGS stability and negative cooperativity [12]. Mutations at these hydrophobic interface residues decrease thermal stability and alter cooperativity, with mutations to Val45 generally producing larger effects than mutations to Val44 [12]. These findings led to the conclusion that Val44 and Val45 lie along the hGS allosteric pathway and that the dimer interface mediates communication between the active sites. The present interface analysis supports that conclusion, as one of the strongest ligand-dependent regions lies within the 42– 46 interface segment. However, the present data also indicates that the interface contribution is distributed rather than confined to Val44 and Val45 alone.

Strong interface interactions cannot be assumed to also play a role in allosteric communication. Previous studies of the electrostatic interface residues Asp24, Ser42, and Arg221 show that alanine mutations at these sites decrease activity, alter substrate affinity, and reduce thermal stability [10]. However, those mutations have little effect on the Hill coefficient [10]. Thus, strong electrostatic interactions across the hGS dimer interface are essential for activity and stability, but they do not appear to be principally responsible for negative cooperativity in hGS. This distinction is useful for interpreting the present analysis of the dimer interface. Some ligand-dependent interface contacts affect dimer stability or packing, whereas others participate more directly in allosteric communication. The present results, therefore, support a model in which the interface acts as a structural crossing point for allosteric communication between distant active sites.

WISP pathway analysis further supports the proposed view of allosteric communication. Most residues that participate in transmission are shared between the product- and reactant-bound states, with 64.1% of residues common to both pathways. However, the pathways are not identical: 30.8% of transmission residues are specific to the product pathway, whereas 5.1% are specific to the reactant pathway. Thus, product- and reactant-bound hGS appear to use a largely conserved communication scaffold that is modulated by ligand identity. This distinction between shared and ligand-specific pathway residues may be broadly important for understanding enzyme allostery because it suggests that different ligand states do not necessarily require entirely separate communication routes. Instead, ligand identity may redistribute the strength, direction, or functional consequence of communication within a common allosteric motif. These findings are consistent with the previous theory that hGS allostery arises from a cumulative effect of many weak interactions rather than one or two strong inter-subunit bonds. In this model, strong electrostatic contacts maintain the dimeric enzyme in an active structural state, while weaker hydrophobic and loop-mediated interactions provide the flexibility required for ligand-dependent communication.

The present findings also agree with prior work on the active-site proximal S-loop. The S-loop forms part of the γ-GC binding site and contains residues that directly affect substrate binding and allosteric behavior. Previous studies show Arg267 and Tyr270 bind γ-GC, that patient mutations occur at these positions, and that experimental mutations in the S-loop can reduce activity and diminish cooperativity [8] [33] [34]. For example, R267K and P272A are reported to reduce cooperativity, whereas other S-loop mutations alter substrate binding with smaller effects on cooperativity [8]. These observations indicate that allosteric communication in hGS begins with substrate-specific binding interactions at the active site. The present simulations are consistent with this idea, as ligand identity at one site is associated with structural changes at the empty partner site, suggesting that the substrate-binding loops also serve as the input region for inter-site communication, in addition to acting as local catalytic elements.

Prior G-loop and A-loop findings provide complementary insights into the roles of these loops in communicating cooperativity. Previous studies of the G-loop glycine triad show that Gly369 and Gly370 are essential for activity, active-site closure, and positioning ATP’s γ-phosphate [11]. Mutations at Gly369 and Gly370 strongly reduce activity and impair ligand binding and loop closure [11]. Similarly, studies of Asp458 in the A-loop showed that D458A, D458N, and D458R mutants retain overall stability but lose negative cooperativity and show large changes in substrate binding, especially for glycine [9]. These findings are important because they show that hGS allostery can be altered by local loop changes that do not grossly alter the enzyme’s global conformation. The present empty-site volume and hydration results fit this pattern by suggesting that ligand binding changes the structural readiness of the empty partner site through coupled loop and interface motions rather than by disrupting the global structure.

The involvement of the dimer interface 212–236 α-helical region in the presently proposed allosteric pathway is also corroborated by previous work on γ-GC binding. Residues in this region, including Glu214, Asn216, Gln220, and Arg221, are implicated in γ-GC binding, active-site organization, dimer-interface stability, and previously proposed communication routes [4] [10]. The present pathway and interface analyses identify the 212–236 region as a major component of the ligand-dependent response, suggesting this region serves as a bridge between substrate-proximal contacts and the dimer interface. It is therefore reasonable to consider the 212–236 segment as part of the broader structural pathway connecting ligand binding to inter-subunit communication.

The presently proposed pathway also has implications for interpreting hGS variants outside the catalytic center. The shared pathway includes residues Leu27, Leu32, Arg34, Ser42, Ile217, and Phe218 of both subunits, placing the conserved communication route near two mutation-sensitive structural regions: the 42–46 dimer-interface segment and the 212–236 helical/interface region. Ser42 directly participates in the shared allosteric pathway, and prior work shows that Ser42Ala decreases thermal stability while preserving negative cooperativity, indicating that this position contributes strongly to dimer stability but may not be required for allosteric communication [10]. Residue Ser42 also lies adjacent to Val44 and Val45, which have clearer evidence for involvement in the allosteric pathway [12]. Similarly, residues Ile217 and Phe218 lie near Asp219 and Arg221, positions associated with altered enzyme behavior and disease-linked regions. These overlaps suggest that the pathway identified here may help explain how mutations outside the catalytic center can impair hGS activity, stability, or inter-subunit communication.

The clinical relevance of this model is further reinforced by observed patient mutations at Tyr270 and Arg267, which are associated with low glutathione levels, hemolytic anemia, metabolic acidosis, neurological symptoms, and, in severe cases, premature death [33] [34] [8]. Prior experimental and computational work indicates these effects alter γ-GC binding, disrupt the active-site structure, and interfere with allosteric communication [8]. The present results extend that interpretation to the partner active site and the dimer interface, suggesting that engineered or pathogenic mutations impair hGS by altering the communication network that coordinates the two sites of the homodimer in addition to disrupting catalysis at the active site.

The present findings may also inform the study of cooperativity in other oligomeric enzymes. In many multimeric enzymes, active sites are separated by substantial distances, making it difficult to explain cooperative behavior at the atomic level. This model of hGS allostery suggests that negative cooperativity can arise when ligand binding at one site alters the ensemble of local conformations explored by a distant partner site. Importantly, this effect can be detected by changes in active-site metrics such as hydration, pocket volume, loop positioning, and distributed residue interactions, even when large global structural transitions are not apparent. Thus, the present study provides a starting point for a generalizable approach to identifying how distant active sites influence one another. This framework may be useful for other enzymes in which mutations outside known functional hotspots alter activity, stability, or cooperativity.

Although the present findings support a model of distributed inter-site coupling, alternative interpretations are considered. While the larger, less-hydrated empty partner site in the product-bound state may reflect allosteric remodeling, it may also be influenced by how the active-site residues are defined, by local pocket breathing, or by water exchange. Similarly, analysis of the dimer interface identifies residues whose interactions change with ligand state, but it does not prove each contact is required for communication. This limitation is worth considering, as prior Asp24/Ser42/Arg221 studies show large effects on stability and activity do not necessarily imply a direct role in negative cooperativity. Thus, the present interface analysis should be interpreted as a means of identifying candidate structural participants in ligand-dependent remodeling without assigning causality to every implicated residue.

These limitations, however, do not eliminate the central interpretation of the present research, as several independent readouts all converge to the same conclusion. Empty-site volume, empty-site hydration, interface interactions, and WISP pathways all indicate that ligand identity in one subunit affects the partner subunit. Prior mutagenesis studies of Val44/Val45, the S-loop, the G-loop, and Asp458 further demonstrate that hGS allostery depends on both substrate-proximal loop interactions and the dimer interface architecture [12] [8] [11] [9]. The present study links these previously studied elements into a single structural model: ligand binding is sensed at the active-site loops, transmitted through a shared ligand-modulated residue network, and expressed as a remodeled partner-active-site.

Many aspects of this pathway warrant further study. Mutagenesis of shared pathway residues, particularly previously unstudied residues in the 42–46 loop, the S-loop, and the 212–236 region, would help determine whether these positions are required for cooperativity, stability, or substrate binding. Such experiments should distinguish between residues that stabilize the dimer and those that transmit allosteric communication. For example, mutations adjacent to Val44/Val45 would be expected to test the hydrophobic interface pathway, whereas mutations adjacent to Asp24/Ser42/Arg221 would test whether altered contact rewiring reflects communication, stability, or both. Mutations in the Arg267, Glu214, Asn216, Gln220, Ile217, and Phe218 regions would test the connection between γ-GC binding and the currently proposed communication scaffold.

Further kinetic work should determine whether these mutations alter the Hill coefficient, substrate affinity, catalytic rate, or all three. Structural and biophysical approaches, including HDX-MS, NMR, cryo-EM, crosslinking, circular dichroism, and differential scanning calorimetry, could test whether the predicted interface and partner-site changes occur experimentally. It would be especially useful to determine whether mutations that diminish negative cooperativity also alter the empty-site volume and hydration effects observed here. The present results provide a useful framework for such studies. More broadly, this work shows how negative cooperativity can be understood as an emergent property of ligand-sensitive structural networks that connect distant active sites. By linking substrate identity, interface remodeling, pathway residues, and partner-site effects into a single model, hGS provides a mechanistic example of how oligomeric enzymes can fine-tune activity through distributed communication.

## Conclusions

In conclusion, this study establishes a new atomistic model for understanding negative cooperativity in human glutathione synthetase. This is the first study to define how reactant- and product-bound states of hGS remodel the empty partner active site differently, and it is the first to map the inter-subunit communication network linking ligand binding in one subunit to structural changes in the other. By integrating microsecond-scale molecular dynamics simulations, empty-site structural analysis, interface contact mapping, and Weighted Implementation of Sub-optimal Path (WISP)-based network analysis, this work provides a residue-level description of long-range (∼40 Å) coupling across the hGS homodimer.

The research reported herein supports a model in which hGS negative cooperativity arises from ligand-dependent remodeling of a shared inter-subunit communication network. Binding of reactants or products to one subunit induces distinct structural states in the empty partner site, as evidenced by changes in empty-site volume, hydration, interfacial contacts, and WISP-derived communication pathways. These effects are not readily explained by a single local contact or by the simple opening and closing of the active site. Rather, they show that hGS uses a shared distributed communication scaffold that links distant active sites through coordinated changes in active-site loops, interface contacts, and pathway residues. This provides a mechanistic explanation for how two active sites separated by ∼40 Å can remain functionally coupled.

The model developed in this research both confirms and extends prior structural and biochemical studies of hGS. Active-site loops, including the S-loop, G-loop, and A-loop, function as input regions for allosteric communication in addition to acting as local catalytic elements. The dimer interface serves as a structural crossing point between monomer subunits, with regions such as the 42–46 segment and the 212–236 region helping couple substrate-proximal interactions from one active site to the partner active site. Strong electrostatic interface contacts are proposed to maintain dimer stability and activity, whereas weaker hydrophobic and loop-mediated interactions provide the flexibility required for ligand-sensitive communication.

A major outcome of this work is that it converts the broad concept of hGS negative cooperativity into specific, testable structural hypotheses. Although the present simulations do not establish the causal role of every residue in the proposed allosteric pathway, they identify defined regions for experimental validation. Residues near the 42–46 interface segment, the S-loop, the 212–236 region, and shared pathway positions such as Leu27, Leu32, Arg34, Ser42, Ile217, and Phe218 provide useful targets for future mutagenesis, kinetic analysis, and structural validation. Experiments that distinguish effects on stability, substrate binding, catalytic rate, and Hill coefficient are especially important for separating structural support from allosteric transmission.

Overall, this study advances understanding of hGS negative cooperativity from a phenomenological observation to a mechanistic, residue-level model of inter-subunit communication. Ligand binding at one active site alters the structural readiness of the distant partner site by redistributing interactions across a shared, ligand-sensitive communication scaffold. This mechanism provides a framework for interpreting how mutations outside of known functional hot spots can affect hGS activity and cooperativity. More broadly, the present research illustrates how oligomeric enzymes can coordinate the reactivity of distant active sites through distributed structural networks, and it provides a computational strategy for identifying experimentally testable allosteric pathways in multimeric enzymes.

## Acknowledgements

During manuscript preparation, the authors used AI assistance from Grammarly and OpenAI for language polishing, passive-to-active voice restructuring, early feedback on drafts, title suggestions, code completion, debugging, and code refinement. All AI-assisted outputs were critically reviewed, edited, tested, and validated by the authors, who take full responsibility for the final manuscript, analyses, and conclusions.

## Footnotes

† Glu, Lys, Glu, Arg, Asn, Ile, Phe, Asp, Gln, Arg, Ala, Ile, Glu, Asn, Glu, Leu, Leu, Ala, Arg, Asn, Ile, His, Val, Ile, Arg

